# Cargo competition for a dimerization interface stabilizes a protease adaptor in *Caulobacter crescentus*

**DOI:** 10.1101/2020.06.12.148478

**Authors:** Nathan J. Kuhlmann, Dylan Doxsey, Peter Chien

## Abstract

Bacterial protein degradation is a regulated process aided by protease adaptors that alter specificity of energy dependent proteases. In *Caulobacter crescentus*, cell-cycle dependent protein degradation depends on a hierarchy of adaptors, such as the dimeric RcdA adaptor which binds multiple cargo and delivers substrates to the ClpXP protease. RcdA itself is degraded in the absence of cargo and how RcdA recognizes its targets is unknown. Here we show that RcdA dimerization and cargo binding compete for a common interface. Cargo binding separates RcdA dimers and a monomeric variant of RcdA fails to be degraded, suggesting that RcdA degradation is a result of self-delivery. Based on HDX-MS studies showing that different cargo rely on different regions of the dimerization interface, we generate RcdA variants that are selective for specific cargo and show cellular defects consistent with changes in selectivity. Using the same interface for dimerization and cargo binding offers an ability to limit excess protease adaptors by self-degradation, while providing capacity for binding a range of substrates.

**Significance Statement:** Energy-dependent proteases broadly regulate bacterial physiology and development. Adaptor proteins tune the substrate specificity of proteases to only degrade selective substrates during the bacterial life cycle and during times of cellular stress. In the process of delivering cargo to their respective proteases, adaptor proteins are inherently protected from degradation until the delivery is complete. How protease adaptors can recognize a wide range of cargo while maintaining stringent specificity and how this process results in stabilization of adaptors remains unclear. Here, we show that direct competition for distinct regions of the dimer interface of the RcdA adaptor by its cargo protects RcdA from degradation by the ClpXP protease, and that this interface can be selectively perturbed in a rational manner with biochemical and physiological consequences.

**Highlights:**

- Cargo binding of RcdA cargo competes with dimerization
- Dimerization of RcdA is necessary for self-degradation by ClpXP
- RcdA can deliver either cargo or other RcdA subunits to ClpXP
- Different regions of the dimerization interface are needed for different cargo

## Introduction

Controlled protein degradation regulates key physiological processes in all domains of life. In bacteria, AAA^+^ (<u>A</u>TPases with <u>A</u>ssociated <u>A</u>ctivities) proteases control the regulated destruction of misfolded and native substrates to manage cellular stress responses, cell-cycle progression, physiological development, and general protein quality control maintenance (Mahmoud and Chien, 2018). The Hsp100/Clp family of proteases, which includes the AAA+ protease ClpXP, share structural features and are critical for degrading factors to promote normal cell physiology in bacteria and organelles (Truscott et al., 2011, Olivares et al, 2018). These energy-dependent machines recognize substrates using an oligomeric unfoldase which translocate these targets into peptidase chambers that non-specifically cleaves proteins into smaller fragments (Sauer and Baker, 2011).

To ensure that only specific proteins are degraded by the protease complex, bacteria make use of additional accessory factors, called adaptors, that tune the substrate specificity of the protease (Kuhlmann and Chien, 2017). Many adaptors act as scaffolds, tethering specific targets to the protease and increasing local concentration to drive degradation. One such example is the SspB adaptor, which binds and scaffolds ssrA-tagged substrates and the extracytoplasmic stress response factor N-RseA to ClpXP (Levchenko et al., 2000, Chien et al., 2007, Flynn et al., 2004). In *Caulobacter crescentus*, adaptors can work additively to provide increasing levels of substrate specificity to the ClpXP protease (Joshi et al., 2015). In this system, the CpdR adaptor activates ClpX and promotes binding of the RcdA adaptor (Iniesta et al., 2006, Lau et al., 2015; McGrath, et al. 2006). RcdA can bind a third adaptor, PopA, which requires cyclic-di-GMP to promote degradation of CtrA, a major regulator of the *Caulobacter* cell cycle (Quon et al., 1996, Reisenauer et al., 1999, Jenal et al., 1998, Ozaki et al., 2014, Duerig et al., 2009; Smith, et al. 2014). In the absence of PopA, RcdA can also deliver substrates such as the polar cue dependent chromosome segregation protein SpbR (originally annotated as CC2323) (Wang and Bowman, 2019; Joshi et al., 2015) and the stalk synthesis protein TacA (Biondi et al, 2006; Bhat et al., 2013, Joshi et al., 2015).

RcdA was crystallized as a homodimer, where two three-helix bundle subunits dimerize via conserved hydrophobic residues in the second helix (Taylor et al., 2009). The disordered C-terminus is necessary for interactions with CpdR/ClpX, for delivery of all RcdA-dependent cargo, and for self-degradation (Taylor et al., 2009, Joshi et al., 2015, Joshi, et al. 2017). Upon cargo binding, the CpdR-mediated degradation of RcdA is suppressed, but how this self-degradation is regulated is unclear (Joshi., et al 2017). Direct binding between RcdA and its cargo has been shown using purified components (Joshi et al., 2015, Joshi et al., 2017), but the details of how RcdA binds and regulates the turnover of a diverse range of cargo remains unclear.

Here, we show that cargo binding competes with RcdA dimerization by competition for overlapping interfaces. Based on biophysical measurements we determine that while RcdA is a dimer in solution on its own, the adaptor binds cargo as a monomer. We generate a constitutively monomeric variant by mutating the predicted dimer interface and show that RcdA dimerization is required for self-degradation. Interestingly, this variant is deficient in delivering the substrates SpbR and TacA for degradation but facilitates PopA-dependent CtrA degradation as normal. We use hydrogen-deuterium exchange mass spectrometry to map regions of RcdA important for cargo binding and find that different substrates rely on different sites of the dimer interface. Mutations at these regions result in adaptor variants that are defective for degradation of specific substrates and expression of these variants alter cell physiology consistent with this change in specificity. Taken together, our data show how RcdA can deliver either cargo or itself for degradation and how a large interface, normally masked by dimerization, can be used to capture a range of substrates.

## Results

### Cargo Binding Competes with RcdA Dimerization

Previous studies have established RcdA is a homodimeric protein in solution (Taylor et al., 2009) that can directly bind to its cargo (Joshi et al., 2015). We began our studies by exploring RcdA-cargo binding using size-exclusion chromatography with multi-angle light scattering (SEC-MALS) to accurately measure molar mass. SpbR, a RcdA-dependent cargo responsible for inhibiting centromere translocation (Wang et al., 2019) is a 44 kDa monomer in solution, which elutes at a similar time as dimeric RcdA (34 kDa, Figure 1A). The combination of both proteins results in a complex with an experimental mass of 60 kDa (Figure 1A). Our data is consistent with a monomer of RcdA (17 kDa) binding a monomer of SpbR (44 kDa) and inconsistent with a dimer of RcdA binding a monomer of SpbR (a predicted mass of 77 kDa) (Table 1). We confirmed this monomer:monomer stoichiometry by using analytical ultracentrifugation and found a similar mass of 60 kDa for the RcdA:SpbR complex (Supplemental Figure 1A).

**Figure 1:**
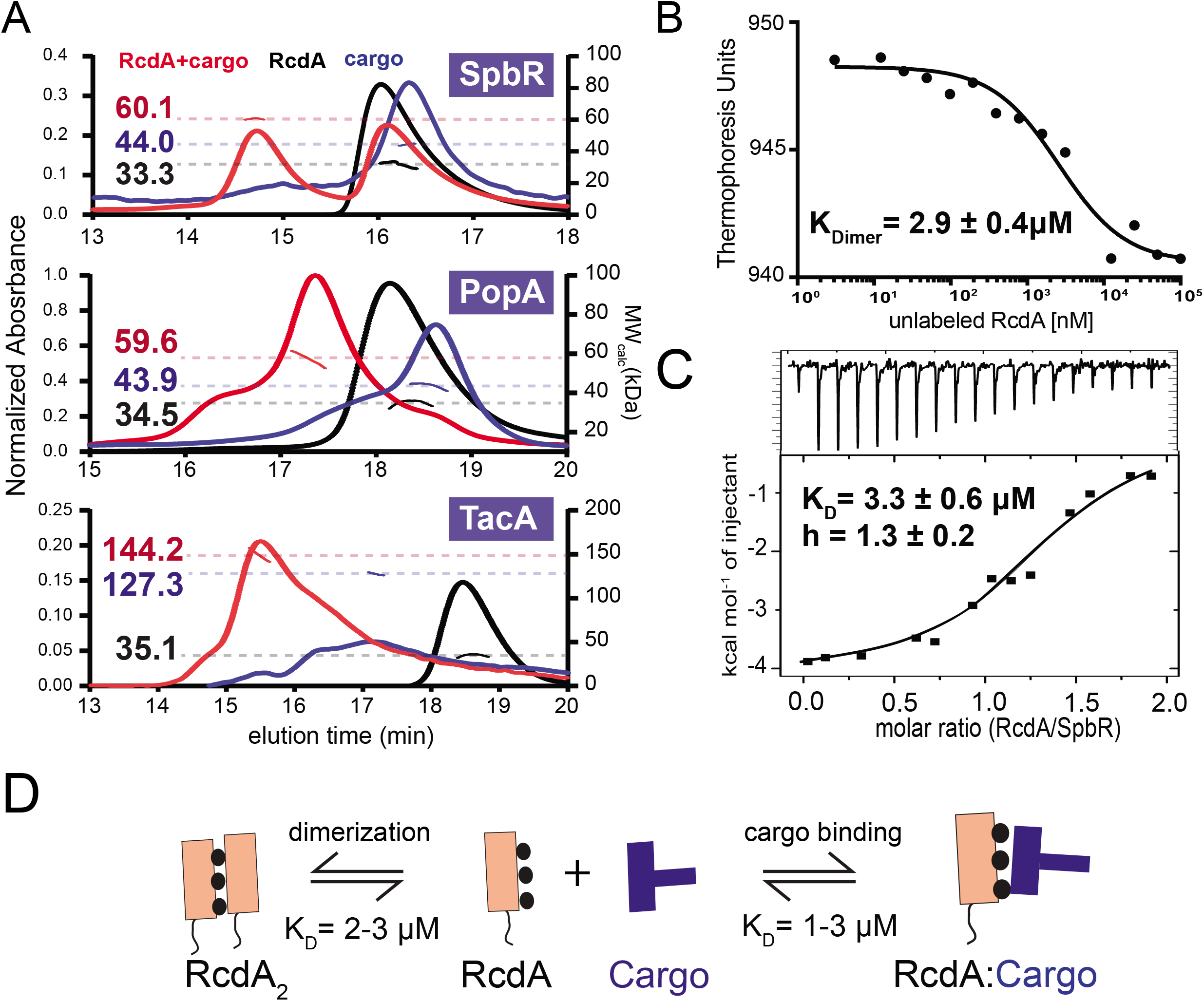
Cargo binding competes with RcdA dimerization. **A)** SEC-MALS traces of different RcdA-cargo complexes. Representative SEC-MALS traces of the RcdA, cargo, and the RcdA-cargo complexes. All SEC-MALS Data is from a representative SEC-MALS experiment that was performed at least twice. **B)** Microscale thermophoresis curve for determining the dimerization constant of RcdA. Data is from a representative MST titration that was performed in triplicate. **C)** ITC thermogram of the SpbR-RcdA complex formation. The sample syringe was loaded with 400μM RcdA and titrated into a cell containing 40μM SpbR. Data is from a single titration. **D)** Cartoon illustration highlighting the competition between dimerization and cargo binding.

**Table 1:** Cargo Binding Competes with RcdA Dimerization. Table of molecular weights of the different RcdA-cargo complexes. The experimental column shows the complex mass measured by SEC-MALS. The stoichiometry column gives the stoichiometry of each protein consistent with the experimental mass. The predicted column lists the EXPASY predicted mass of each stoichiometry listed. Bolded text is the experimentally measured complex masses displayed in Figure 1.

|  | Experimental (kDa) | Stoichiometry | Predicted (kDa) |
| --- | --- | --- | --- |
| RcdA | 33.3 | RcdA<sub>2</sub> (dimer) | 37.0 |
|  |  | RcdA (monomer) | 19.0 |
| SpbR | 44.0 | SpbR (monomer) | 43.4 |
| **RcdA + SpbR** | **60.1** | **RcdA:SpbR** | **62.4** |
| TacA | 127.3 | TacA<sub>2</sub> (dimer) | 127.0 |
| **RcdA + TacA** | **144.0** | **RcdA:TacA<sub>2</sub>** | **146.0** |
| PopA | 43.9 | PopA (monomer) | 47.4 |
| **RcdA + PopA** | **59.6** | **RcdA:PopA** | **66.4** |

We next tested other RcdA cargo. PopA, the third adaptor in the ClpX adaptor system, runs as a 43 kDa monomer (Figure 1A). In the presence of RcdA, the complex mass is 59 kDa consistent with a monomer of PopA (43 kDa) binding a monomer of RcdA (17 kDa) (Figure 1A; Table 1). TacA, a transcription factor involved in stalk biogenesis, shows a molar mass consistent with a 127 kDa dimer. When bound to RcdA, the complex runs at an apparent mass of 144 kDa, again consistent with a monomer of RcdA binding to a dimer of TacA (Figure 1A; Table 1). Taken together, these data suggest that the active stoichiometry of RcdA that binds cargo is as a monomer, and that binding of cargo competes with dimerization (Figure 1D).

The binding constant between RcdA and SpbR was measured by isothermal calorimetry (K_D_ = 3.3 μM). This was comparable to the dimerization constant of RcdA subunits as measured by microscale thermophoresis (K_D_ = 2.9 μM) (Figures 1 B,C) using fluorescein-labeled RcdA. Additional measurements of SpbR and PopA binding using fluorescence polarization confirmed that dimerization and cargo binding were similar in strength (Supplemental Figure 1C-E).

### RcdA Dimerization is Necessary for its own Degradation

Previous studies have shown that cargo binding inhibits the self-degradation of RcdA by the ClpXP protease (as shown with a SpbR variant cargo in Figure 2B) (Joshi, et al. 2017), a process that relies on the C-terminal tail of RcdA acting as both a degron and a tether to ClpX (Joshi, et al 2017). Originally, we speculated that a stabilizing conformational change occurred upon cargo binding that prevents adaptor recognition by ClpXP. Since cargo binding competes with RcdA dimerization (Figure 1), we considered a simpler model where a monomer of RcdA binds to a CpdR-activated ClpX and tethers a bound cargo. In this model, adaptor monomers can deliver substrates such as TacA, SpbR or another RcdA monomer - or can recruit additional factors like PopA. If this model is correct, then robust RcdA degradation would require a homodimer containing two intact C-terminal tails (Figure 2A). We tested this by making mixed heterodimers of a full-length RcdA and an excess of a truncated, non-degradable RcdAΔC variant (Joshi, et al. 2017). We hypothesized that in this heterocomplex, the full-length monomer would anchor to CpdR-ClpX, but not be degraded because it was not presented as a cargo (Figure 2A). Indeed, loss of the full length RcdA in the heterodimer was much slower when compared to the homodimer (Figure 2B). This is consistent with our model that RcdA delivers itself as a homodimer that requires two C-terminal degron tails in both subunits, with the slow exchange of heterodimers to homodimers contributing to the slow degradation seen here.

**Figure 2:**
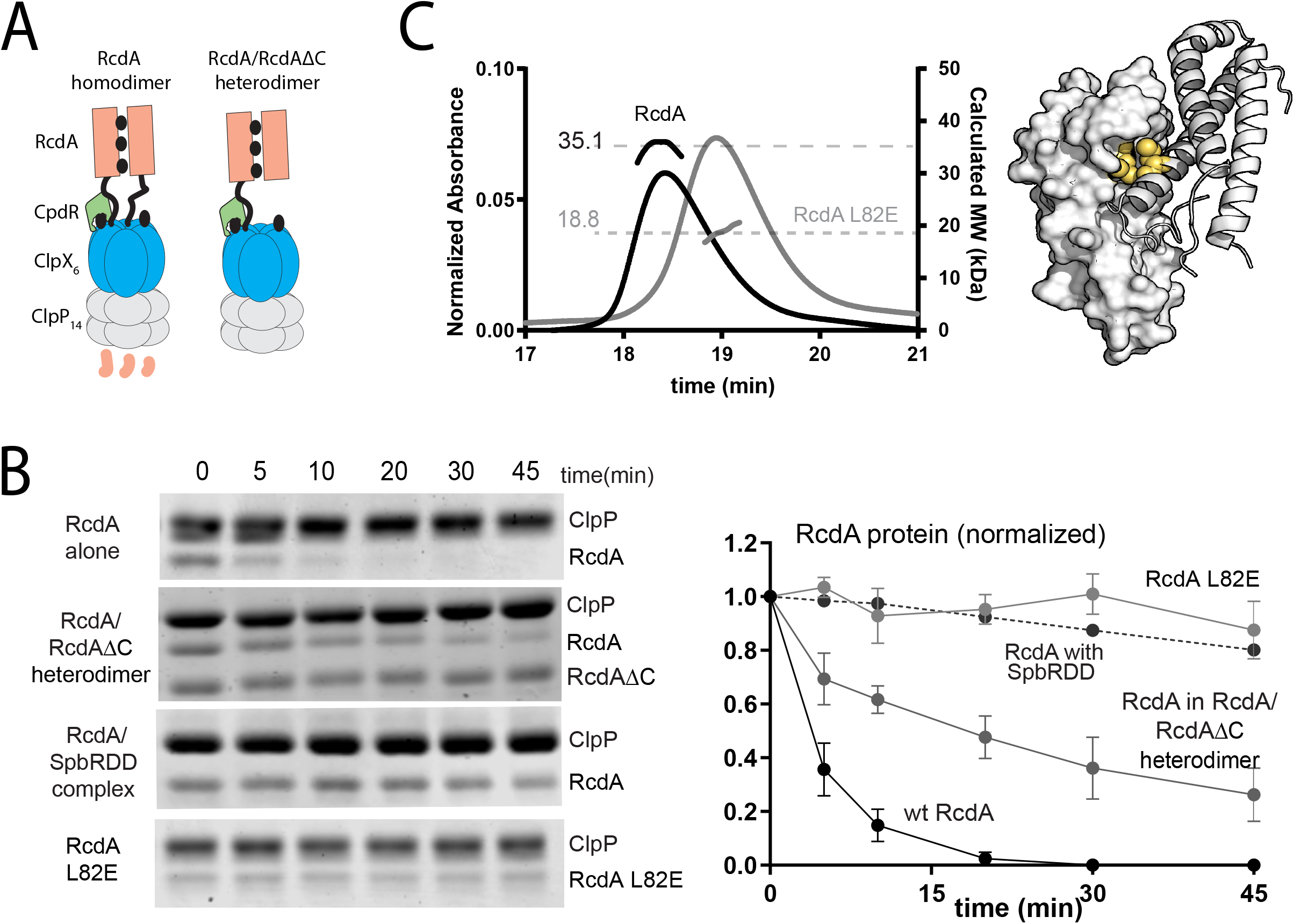
Dimerization is necessary for RcdA degradation. **A)** Cartoon illustration of the proposed degradation model of RcdA. **B)** *In vitro* gel-based degradation of 3 μM RcdA alone, or in the presence of 10uM RcdAΔC, 3μM SpbRDD. Also shown is the degradation of 3 μM RcdA L82E. Quantification of three independent replicates is shown on the right. Coomassie-stained gels were visualized using an Odyssey imaging system (LICOR) for better visualization of the RcdA band. See Supplemental Figure 2 for full gels. **C)** SEC-MALS trace of 25 μM RcdA and 25 μM RcdA L82E. Illustration of the L82 residue (in yellow) in the RcdA (PDB ID: 3CTW) crystal structure where one monomer is surface rendered and the other is in cartoon..

To more completely test our hypothesis that RcdA is degraded as a homodimer, we mutated Leucine 82 in the RcdA dimer interface (as determined from the reported crystal structure (PDB: 3CTW; Taylor et al., 2009)) to a glutamic acid (Figure 2C). In contrast to the wildtype dimer mass (35 kDa), RcdA-L82E behaved as a 19 kDa monomer (Figure 2C) and failed to form dimers with itself or with wildtype RcdA by fluorescence polarization (Supplemental Figure 2C). The monomeric L82E variant was also completely stable *in vitro* in the presence of ClpXP and CpdR (Figure 2B). Finally, while wildtype RcdA is degraded *in vivo*, RcdA L82E is not (Supplemental Figure 2E). These data support a model where dimerization of RcdA is necessary for its self-delivery to the ClpXP protease for degradation.

### The L82E Variant is Defective in Some Cargo Binding and Delivery

We next tested the ability of RcdA L82E to bind and deliver different RcdA cargo. SpbR and TacA are substrates of ClpXP that require RcdA for efficient degradation, while PopA is an adaptor that binds RcdA to expand its substrate profile (Joshi et al, 2015). We used wildtype RcdA or RcdA L82E labeled with fluorescein and fluorescence polarization as a proxy for cargo binding. Using this assay, we found that wildtype RcdA bound all three cargo (n.b., ^DBD^TacA is the minimal domain of TacA needed for RcdA degradation) (Figure 3A; Supplemental Figures 1C and 3A). The monomeric RcdA L82E was unable to form a complex with SpbR and ^DBD^TacA, but surprisingly, bound PopA with affinity equivalent to wildtype RcdA (Figure 3A, Supplemental Figure 3A). SEC-MALS confirmed these changes in cargo binding (Figure 3B, Supplemental Figure 3E).

**Figure 3:**
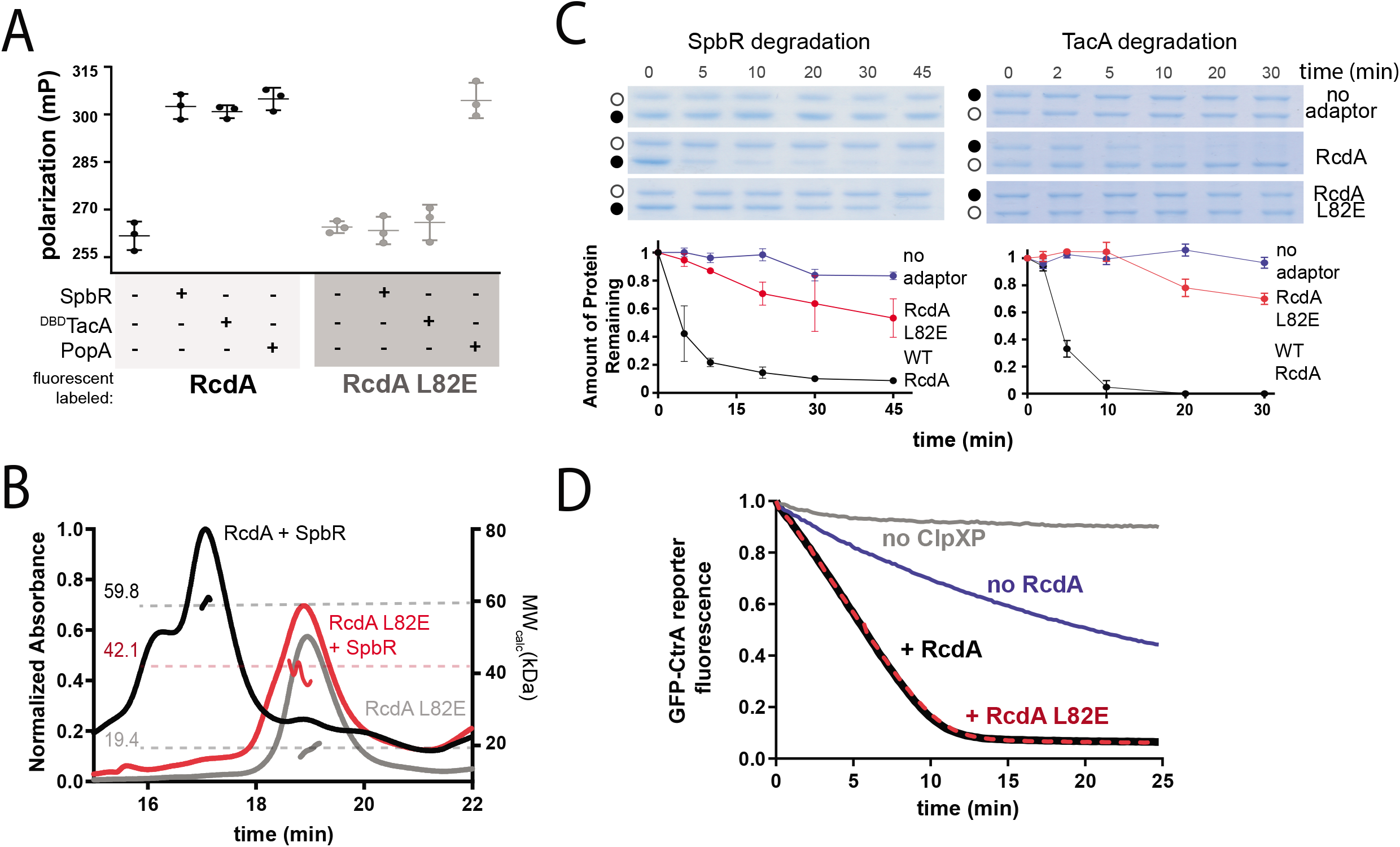
The L82E variant shows differences in cargo binding and RcdA activity. **A)** Fluorescence polarization of labeled RcdA (black) or RcdA L82E (grey) in the presence of SpbR, ^DBD^TacA and PopA. 200nM of each labeled reporter was incubated with 20μM cargo for 45 minutes and the polarization of labeled RcdA or L82E was measured. **B)** Representative SEC-MALS chromatogram of L82E in the presence of SpbR. Native RcdA incubated with SpbR is shown for comparison. **C)** *In vitro* gel-based degradation assay of SpbR or in the presence of RcdA or RcdA L82E. Black dots indicate substrate of interest and white dots indicate the ClpX loading control. Full gels are shown in Supplemental Figure 3. Quantification of three independent replicates is shown below with the mean and error bars (standard deviation). **D)** *In vitro* fluorescence degradation assay of GFP-CtrARD+15 in the presence of RcdA or RcdA L82E. Each fluorescence trace is the average of three independent replicates.

Consistent with defects in SpbR and TacA binding, the L82E variant had a decreased ability to stimulate the CpdR-ClpXP mediated degradation of these substrates *in vitro* (Figure 3C). PopA binds RcdA directly, and in a c-di-GMP dependent manner promotes degradation of CtrA (Smith et al., 2014, Duerig et al., 2009). To test the effects on PopA-mediated substrate delivery, we used a GFP reporter fused to the minimal domains of CtrA needed for regulated degradation (GFP-CtrARD+15; Smith et al., 2014). Unlike the case for TacA or SpbR, the RcdA L82E variant was fully capable of stimulating PopA-mediated CtrA degradation to the same extent as native RcdA (Figure 3D, Supplemental Figure 3D). These data show that disrupting the dimer interface of RcdA reduces some, but not all, substrate binding and delivery. Taken together with the observation that all cargoes can compete with RcdA dimerization, our emerging model is that different substrates rely on different regions of the RcdA dimer interface.

### The dimer interface is protected from exchange in the presence of cargo

To determine how different cargoes interact with RcdA, we used hydrogen deuterium exchange-mass spectrometry (HDX-MS) which measures differences in deuterium uptake in the peptide backbone to determine protein-protein interaction surfaces (Konermann et al., 2011, Chalmers et al., 2011). We first compared high and low concentrations of RcdA to map the dimer interface (Supplemental Figure 4A) and found these data to be consistent with the comparison between monomeric RcdA L82E and the dimeric wildtype RcdA that also highlights the dimerization interface (Figure 4A). We then measured deuterium uptake of RcdA incubated with excess ^DBD^TacA or PopA to define potential interaction surfaces with these cargoes (Figure 4A). Each condition is summarized in a differential deuterium uptake plot as a heat map showing percent deuterium uptake for each peptide and regions of largest protection (>15%) mapped onto a surface rendition of a RcdA monomer (Figure 4A). Individual deuterium uptake plots for selected regions of largest protection are provided in Figure 4B and Supplemental Figure 4C.

**Figure 4:**
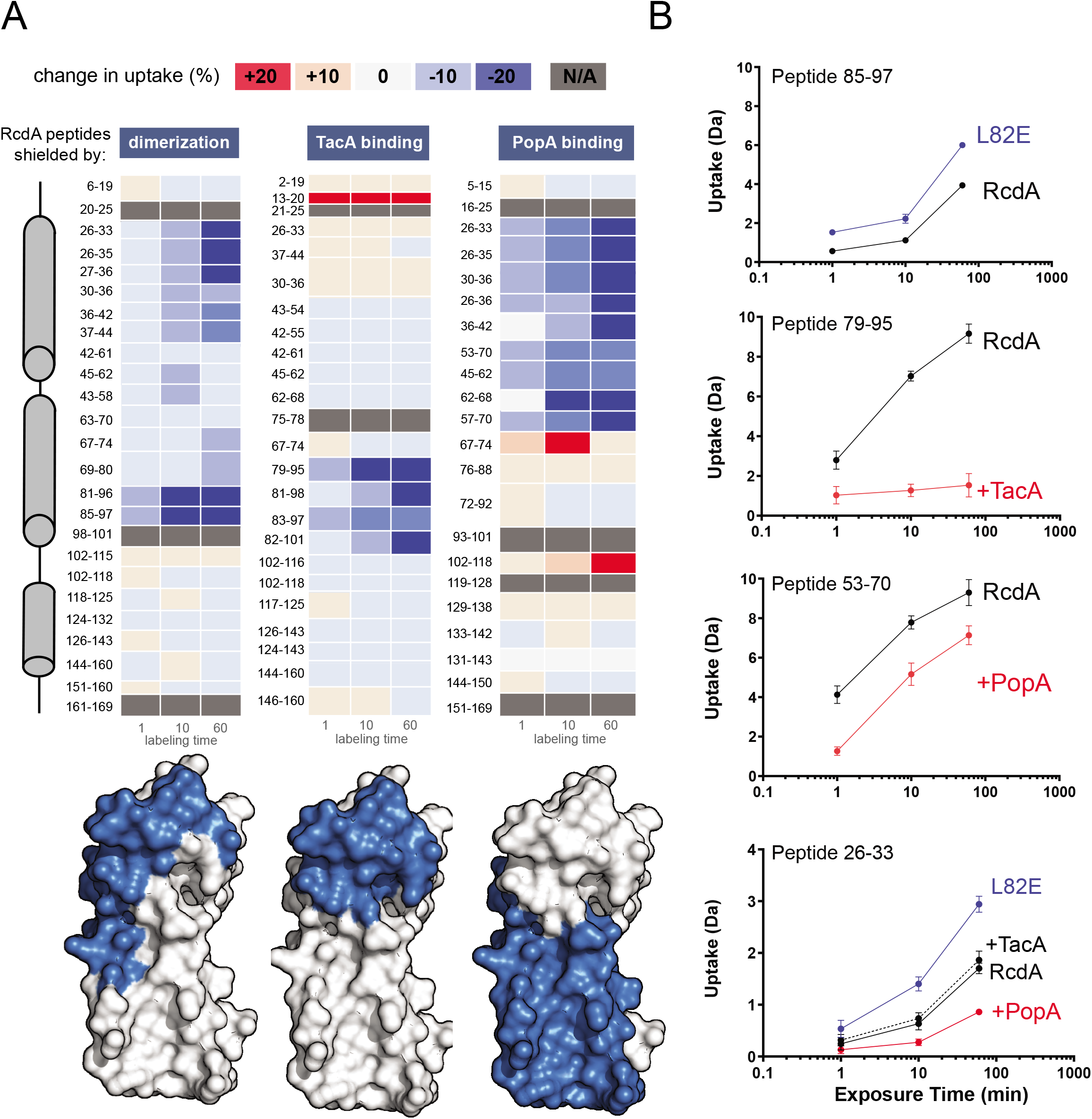
The dimerization interface is protected from deuterium exchange in the presence of cargo. **A)** Differential uptake heatmap plots of 15μM RcdA compared with 15μM RcdA L82E (dimerization), 600nM RcdA incubated with 2OμM ^DBD^TacA (TacA), or 600nM RcdA incubated with 20μM PopA (PopA). Regions of 15% or greater protection are highlighted in blue on a surface rendition of the RcdA monomer structure (PDB ID: 3CTW). **B)** Deuterium uptake plots of selected peptides. Data is plotted as the mean of three replicates with the standard deviation plotted as error bars. See Supplemental Figure 4 for additional peptides.

Based on the PDBePISA server and the 3CTW crystal structure of RcdA, the most buried residues in the dimer interface lie between residues 34-45 and 71-101 (Supplemental Figure 4B). We validated this in solution with our HDX-MS data which highlights residues 26-44 and 80-97 as being most protected in the wildtype dimer (Figure 4A). This result further confirms how the L82E mutation disrupts the dimer interface to generate a monomeric variant. Consistent with our biochemical results in Figure 3, ^DBD^TacA also binds the region of the dimer interface containing the L82 residue with most protection at residues 79-101 supporting our observation that RcdA L82E fails to deliver TacA for degradation. Interestingly, PopA also binds the dimer interface, but principally protects residues 26-70, with the L82 containing region of RcdA showing no substantial protection. We conclude that consistent with our biochemical data, TacA and PopA both bind the RcdA homodimer interface to disrupt dimerization, but they bind at different regions of this interface.

### Mutations in the PopA interaction region highlight differences in cargo binding

Our HDX-MS data highlights two regions in RcdA as protected upon PopA binding, one of which includes a cluster of highly conserved basic residues (R49, K53, R57) (Supplemental Figure 5A). We mutated these residues to glutamic acid to generate a variant that we refer to as RcdA 3E for brevity (Figure 5A). Based on our biochemical studies (Figure 3) we predict that both TacA and SpbR bind similar regions of RcdA, including L82, that are distinct from the regions preferred by PopA (Figure 4). Consistent with this hypothesis, while the RcdA 3E fails to bind PopA based on SEC-MALS, it forms a complex with SpbR with a mass like that of the native complex (Figure 5B). RcdA 3E was unable to deliver CtrA for degradation (Figure 5C), but was active for SpbR and TacA degradation (Figure 5D). These data suggest that while the homodimer interface of RcdA is responsible for cargo binding, PopA and SpbR/TacA rely on different aspects of this interface for effective binding.

**Figure 5:**
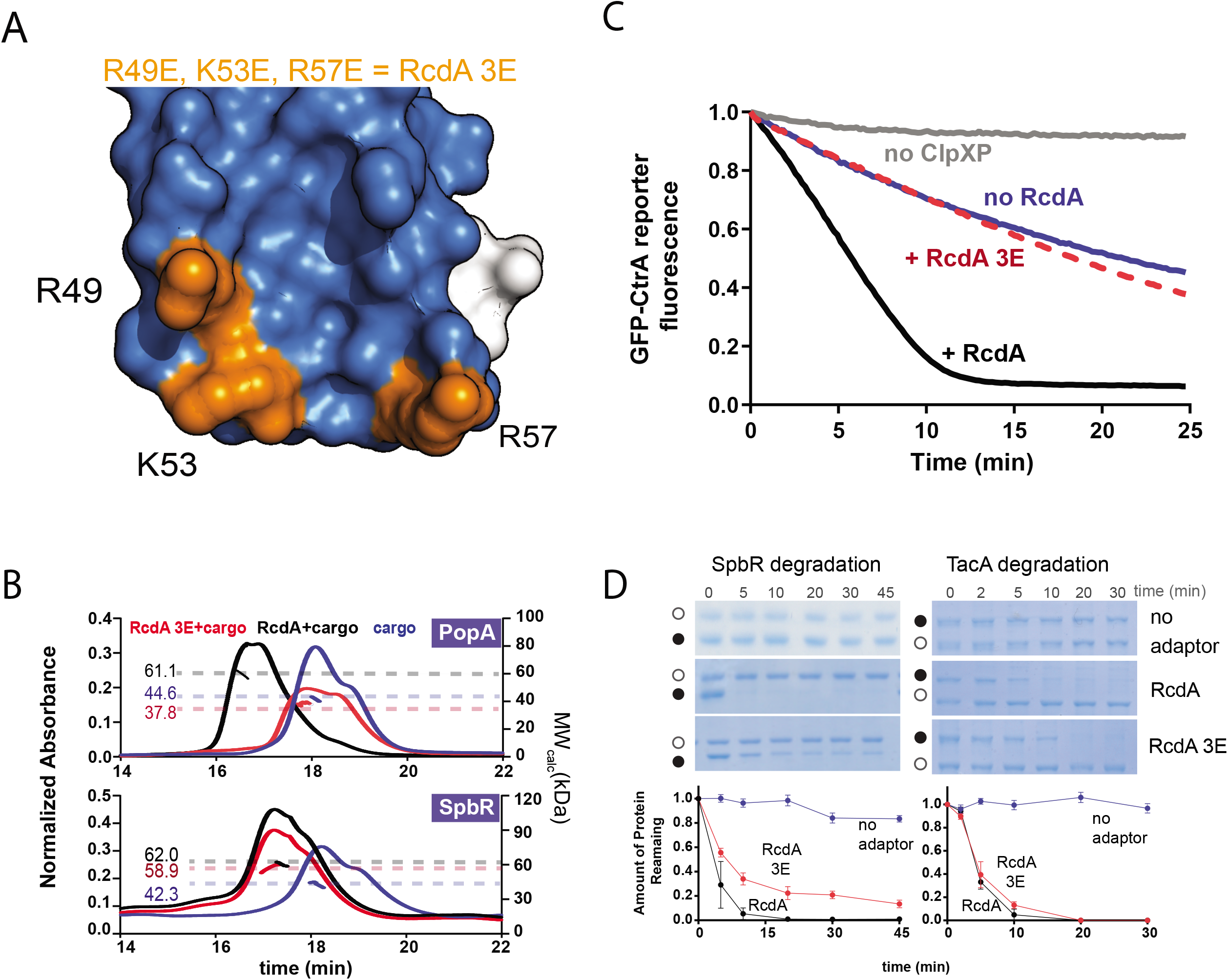
Mutations in the extended surface highlight differences in cargo selectivity and RcdA activity. **A)** Illustration of region of RcdA protected upon PopA incubation (in blue). In orange are the 3 residues (R49, K53, R57) mutated to glutamic acid in RcdA 3E. **B)** Representative SEC-MALS traces for the PopA and SpbR bound-RcdA 3E complexes (red). The complex between native RcdA and PopA and SpbR (black) and the cargo alone (blue) is shown for comparison. **C)** *In vitro* fluorescence degradation assay of GFP-CtrARD+15 in the presence of RcdA or RcdA 3E. Fluorescence traces are the average of three independent replicates. **D)** *In vitro* gel-based degradation assay of SpbR or TacA in the presence of RcdA or RcdA 3E. Black dots indicate substrate of interest and white dots indicate the ClpX loading control. Quantification of three independent replicates is shown below raw gels. See Supplemental Figure 5 for full gels.

### Disruption of the Cargo interaction Surface in vivo

We finally wanted to investigate the *in vivo* consequences of altering RcdA-cargo interactions. We generated strains that express RcdA, RcdA L82E or RcdA 3E at the *xylX* locus in *ΔrcdA* strains, which allows for titration of RcdA using the inducer xylose. *ΔrcdA* strains have defects in normal stalk formation and growth in low percentage agar, (McGrath et al., 2006; Joshi et al., 2015). At low levels of inducer, RcdA variant levels driven by the xylX promoter were similar to that of the wildtype control (Supplemental Figure 6A), and in these conditions, stalk length was compromised in both RcdA L82E and 3E backgrounds (Figure 6A), while growth was affected in the RcdA 3E mutant (Figure 6B). Curiously, under high induction, where RcdA levels are well above normal, stalk length was reduced to less than wildtype for all the alleles tested (included wildtype RcdA) (Figure 6A).

**Figure 6:**
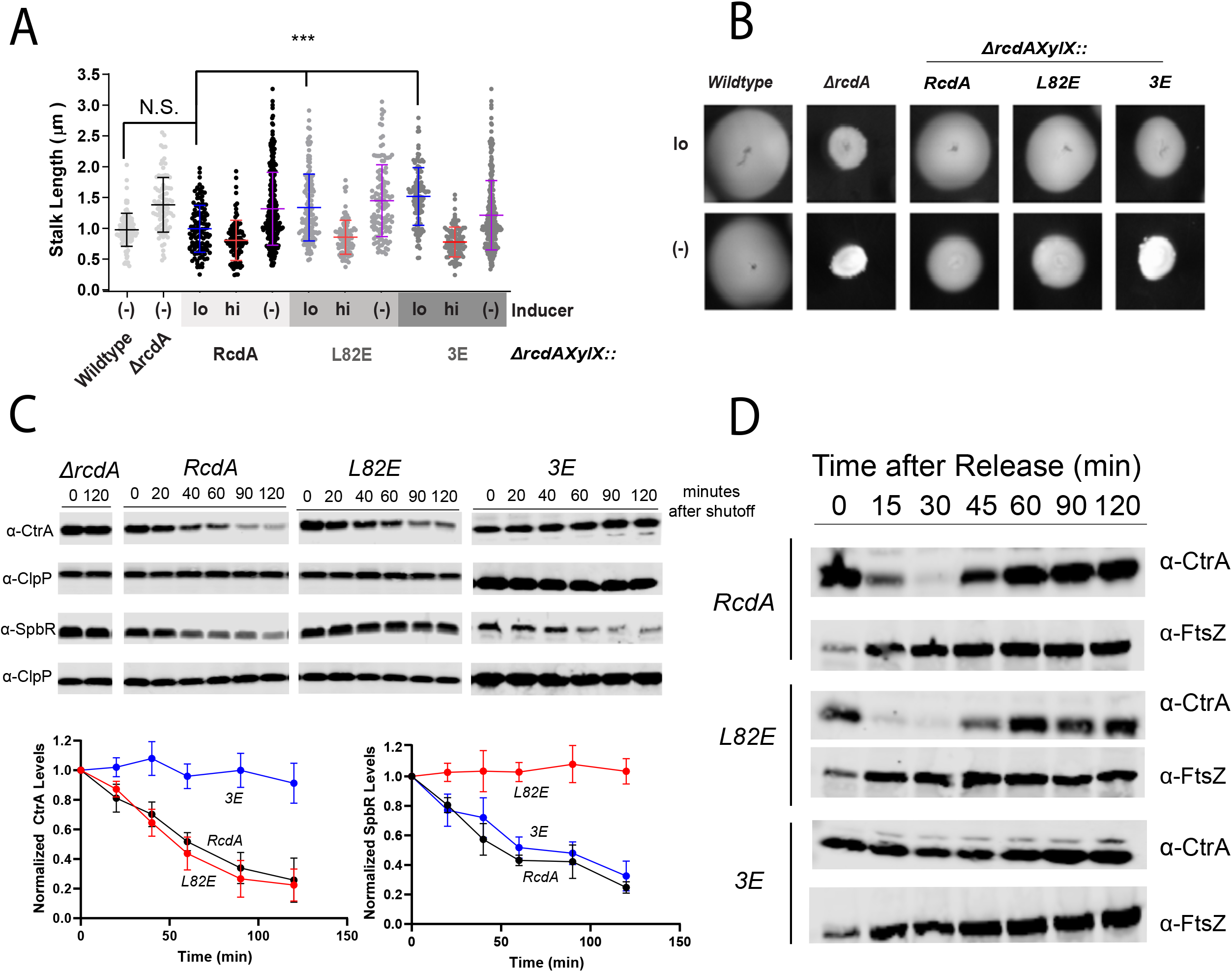
Disruption of the extended RcdA interaction surface affects normal cell physiology. **A)** Stalk length quantification and images of cells expressing RcdA, RcdA L82E or RcdA 3E from the chromosomal xylose locus. Cells grown to exponential phase in media containing 0.2% (hi: high inducer), 0.002% xylose (lo: low inducer) or 0.2% glucose (repressor). Quantification is the average of three biological replicates, with the mean distribution and the standard deviation plotted as error bars. A one-tailed, unpaired t-test with α = 0.05 was used to compare stalk lengths from indicated strains. B) Motility agar assay of each integration strain in PYE + low inducer or repressor agar plates with wildtype RcdA, RcdA(L82E) or RcdA(3E) in each spot. Images have the same absolute perimeter size. **C)** Chloramphenicol shutoff assays monitoring the degradation of SpbR and CtrA in cells expressing wildtype RcdA or RcdA L82E. Quantification of biological replicates (n=3) are shown below. **D)** Synchronized cell growth of each integration strain expressing wildtype RcdA, RcdA L82E or RcdA 3E. Cell samples taken at the indicated time points were blotted against CtrA and FtsZ.

These data suggest that the different substrates stabilized by the different RcdA variants may drive defects in stalk length and agar growth separately. Consistent with this and our *in vitro* results, *in vivo* degradation assays show that strains expressing RcdA L82E are deficient in SpbR degradation (Figure 6C), while strains expressing RcdA 3E are deficient in CtrA degradation during asynchronous growth (Figure 6C) and show defective oscillation of CtrA during the cell cycle (Figure 6D). Taken together with our biochemical results, these data support our model that disrupting dimerization impacts RcdA activity and that separable regions of the dimer interface are important for cargo binding.

## Discussion

Adaptor mediated degradation is critical for bacteria. Our results demonstrate a surprising feature of the cell cycle adaptor system in *Caulobacter*, where binding of a cargo to the RcdA adaptor competes with homodimerization. This competition results in stabilization of RcdA, protecting it from self-degradation by ClpXP, while providing a relatively broad binding surface for cargo binding. We note that the cellular concentration of RcdA estimated by ribosome profiling is 6 μM and SpbR, PopA and TacA are between 2-4 μM each (Aretakis, et al., 2019). This implies that total cargo concentration is likely in excess of RcdA, driving the RcdA equilibrium towards the monomer form and protecting RcdA from degradation until target substrates are delivered by RcdA. Our binding data is consistent with the model that different classes of cargo can interact with specific regions of this surface, with our direct measurement by HDX-MS and our mutation data showing that we can selectively influence particular substrate binding and degradation (Figure 7). Interestingly, CpdR and RcdA are conserved throughout α-proteobacteria, while PopA is only present in *Caulobacter* and closely related bacteria (Brilli, et al, 2010, Ozaki et al., 2014). The region mutated in our RcdA 3E variant may represent the binding interface for currently unknown adaptors that fulfill the role for PopA in other species where CtrA is degraded such as *Sinorhizobium meliloti* (Pini, et al., 2015). Interestingly, recent structures show that the E3 ubiquitin ligase adaptor Skp1 buries its F-box interaction site upon dimerization (Kim, et al. 2020), illustrating that masking of cargo binding sites by self-interactions is more generally found in biological systems.

**Figure 7:**
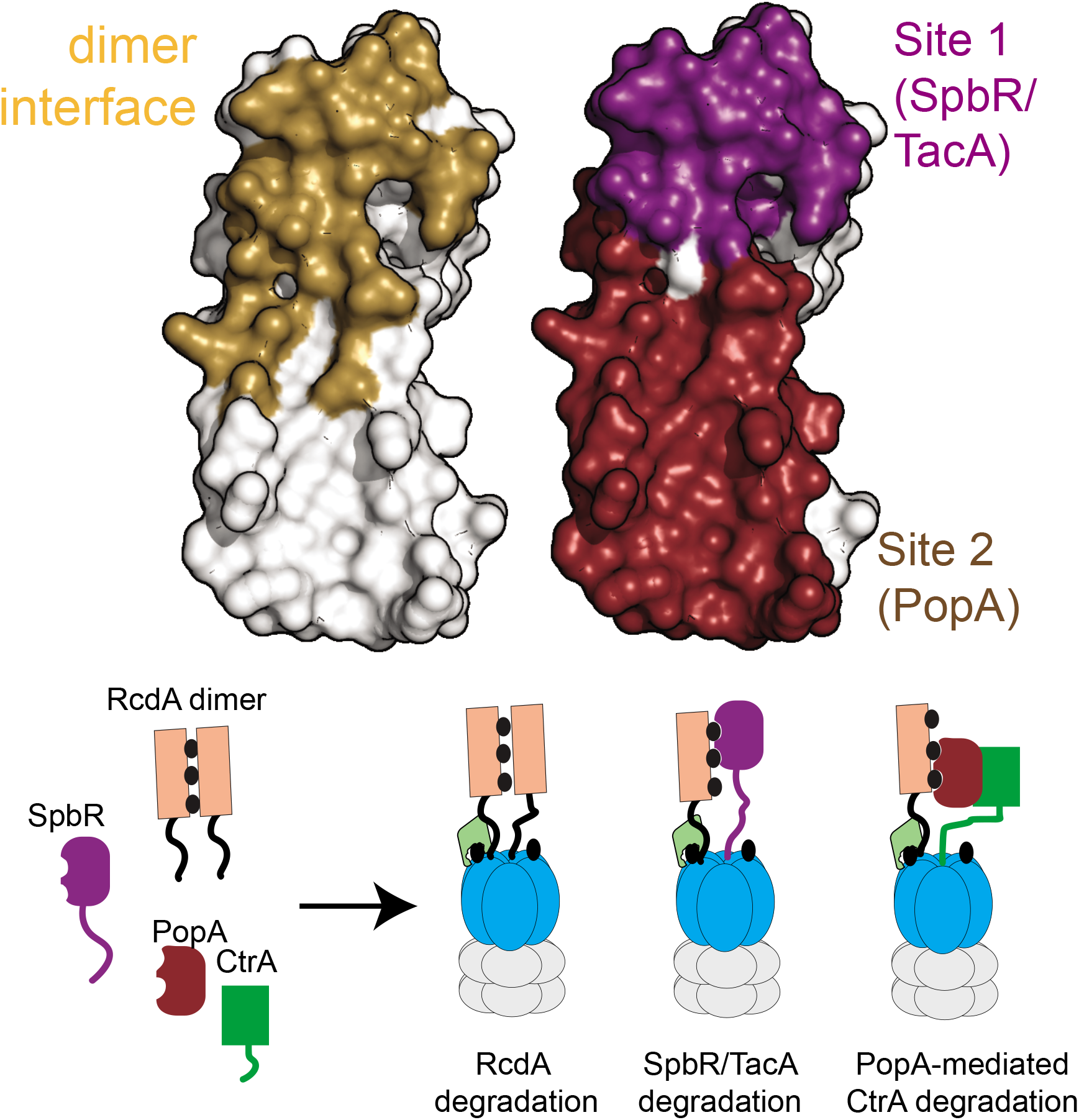
Model of RcdA binding and delivery mechanism. Highlighted structure illustrating the different interactions sites of RcdA. In gold is the dimer interface. In purple is the site involved in the SpbR and TacA interaction, which includes the L82 residue. In ruby is the site involved in the PopA interaction, which includes residues R49, K53, R57. Cartoon model of how the oligomerization state of RcdA affects its activity and self-degradation. When not bound to cargo, RcdA is dimeric and delivered to a CpdR-bound ClpXP for degradation. Cargo binding at different interaction sites by different cargo (SpbR/TacA or PopA) causes dissociation of the RcdA dimer and masks RcdA from its own degradation.

Not all protease adaptors share this mode of binding. The SspB adaptor delivers proteins marked by the ssrA-tagging system to the ClpXP protease (Levchenko et al., 2000, Gottesman et al., 1998). SspB delivery is optimal when two molecules of substrates are bound per SspB dimer (McGuinness et al., 2007, Bolon et al., 2004). Unlike RcdA, SspB binds substrates at sites located at the most distal ends of the dimer, far away from the dimer interface (Levchenko et al., 2003, Chien et al., 2007). Because ssrA-tagged products generally arise from failed translation of damaged or truncated mRNAs, the substrates of SspB are likely incomplete proteins that should be immediately destroyed. Therefore, it is logical that SspB delivery of ssrA-tagged proteins should be optimized for high flux constitutive degradation of targets. By contrast, degradation of substrates by RcdA is a tightly regulated process during the cell-cycle and additional layers of control over delivery of otherwise functional proteins may have useful benefits.

Because protease adaptors catalyze the irreversible destruction of targets, we consider that dimerization of RcdA may limit the potentially toxic overactivation of this adaptor. There are two mechanisms that we think may be at play: First, because the RcdA homodimer is degraded robustly, overlapping the dimer interface and cargo binding site makes levels of RcdA tuned to substrate availability. In the presence of substrates, RcdA monomers are not degraded. Once the substrates are degraded, RcdA homodimers would then be destroyed to limit their activity. Similarly, RcdA would never be fully eliminated as once levels drop below the dimerization constant, the equilibrium shift to the monomeric species would stabilize the adaptor. Second, masking substrate interaction surfaces by dimerization may limit nonspecific binding. Adaptor-substrate affinity can be relatively weak because of the avidity afforded by engaging the AAA+ protease during substrate handoff. By blocking a relatively large binding site with another monomer, nonspecific substrates would be readily competed by RcdA itself, while specific, tighter binding authentic substrates would have longer dwell times on the adaptor and be more likely to be degraded.

## Experimental Methods

### Protein Expression and Purification

BL21(DE3) *pLYS* or X90 cells with expression plasmids for different proteins were grown at 37°C to an OD600 of 0.4-0.6, then induced with 0.4mM IPTG for 3-4 h. Induced cells were then centrifuged at 7,000g for 10 minutes and resuspended in buffer containing 50mM Tris pH 8.0, 300mM NaCl, 10mM imidazole, 10% glycerol, 5mM β-mercaptoethanol and 100mM PMSF and frozen at −80°C until further use. Cells were lysed using a Microfluidizer system (Microfluidics, Newton, MA). The clarified lysate was bound over a Ni-NTA column for affinity purification. H_6_SUMO-tagged proteins were cleaved by Ulp1-his protease (Rood, et al. 2012). Proteins were then purified using size exclusion and anion-exchange chromatography using Sephacryl 200 16/60 and MonoQ 5/50 columns. ClpX and ClpP were purified as outlined in Joshi et al., 2015. Detailed purification protocols are available upon request.

**Cloning and Molecular Biology.** RcdA variants were cloned using around-the-horn site-directed mutagenesis by amplifying the desired plasmid using pET23SUMO-RcdA as a template. To generate the *ΔrcdAXylX::Pxyl* integration strains, the pXMCS-2 was used a template to generate a vector with the RcdA, RcdA(L82E) and RcdA(3E) coding sequences under the xylose promoter. The coding sequence of RcdA, RcdA(L82E) or RcdA(3E) was amplified with complementary overlaps to this vector and the final construct was generated using Gibson assembly method (Gibson et al., 2009). Competent *ΔrcdA Caulobacter*cells were transformed with pXMCS-2-RcdA, RcdA(L82E) or RcdA(3E) containing plasmids and selected on 30μg/μL spectinomycin/streptomycin plates.

### *In Vivo* Protein Stability and Synchrony Assays

Wildtype or *Caulobacter* cells expressing different constructs from a xylose inducible promoter were grown in PYE media with appropriate antibiotic and xylose when required as outlined in the figure legends. Cells were grown to an OD600 of ~0.4 with addition of 0.2-0.002% xylose or glucose. Protein synthesis was blocked by addition of 30μg/mL chloramphenicol and aliquots were taken at the timepoints indicated in the legends. For synchrony experiments, an asynchronous population of cells was grown to an OD600 of ~0.4. Swarmer cells were harvested and isolated using Percoll density gradient centrifugation, then released into fresh PYE media containing 0.002% xylose for progression through the cell cycle.

### Western Blot Analysis

Aliquots withdrawn at indicated time points were spun down, resuspended in 2X SDS sample buffer, boiled at 95 °C for 10 min and then centrifuged. After centrifugation, the clarified supernatant was loaded onto SDS-PAGE gels. Proteins were then transferred onto a nitrocellulose membrane at 20V for 1 hour and probed for monoclonal rabbit anti-RcdA (1:5000), polyclonal rabbit anti-SpbR (1:5000), rabbit anti-FtsZ (1:5000), or rabbit anti-CtrA (1:5000). Following overnight primary probing at 4°C, the membranes were washed 3 times with TBST. Proteins were then visualized using IRdye-labeled goat anti-rabbit antibody (LI-COR Biosciences) at 1:10000 dilution.

### Fluorescence Polarization and Maleimide Labeling

Purified RcdA or RcdA mutants were labeled with Fluorescin-5-Maleimide (Thermo Scientific™). Purified protein at ~8-10 mg/mL was buffer exchanged into labeling buffer (Tris pH 7.0, 150mM NaCL, 2mM TCEP). Fluorescin-5-Maleimide was dissolved in DSMO and added to protein at a 20-fold molar excess to cysteine. Labeling reactions were completed at 4°C overnight. Free dye was removed using a PD-MidiTrap column (GE Healthcare) and Amicon Ultra-15 Centrifugal Filter Units in a buffer containing 20mM HEPES pH 8.5, 100mM KCL, 10mM MgCl2 and 0.05% Tween. Confirmation of protein labeling was verified using a Typhoon imaging system (GE Healthcare). The labeled protein was aliquoted, and flash frozen at −80°C.

Fluorescence polarization binding experiments were initiated by adding 20uL of 200nM labeled RcdA to 20uL of cargo protein at 40μM for final concentrations of 100nM labeled RcdA or L82E and 20μM Cargo. The binding reaction was incubated at 25°C for 1hr. Polarization measurements were read from 40uL of these mixtures using opaque black 384-well plates using a SpectraMax M5 plate reader (Molecular Devices) with excitation and emission wavelengths set at 460 and 540, respectively. Equilibrium binding constants were calculated by fitting the polarization data using GraphPad Prism to a one site, total and nonspecific binding equation P = Pmax~[X]/([X] + K_d_) + NS~[X] + Background, where Pmax is the maximum specific binding value, P is the polarization value, NS is the slope of linear nonspecific binding constrained to be greater than 0, and the background is the polarization value when [X] is 0. Error bars are calculated from the standard deviation between replicates of experiments.

### Isothermal Titration Calorimetry (ITC)

ITC experiments were completed using a Malvern-autoiTC200 automated system (Malvern). Measurements were taken at 25°C. The reference cell was filled with 20 mM HEPES pH 8.5, 100mM KCL and 10% glycerol. The sample cell was loaded with 400μL of 40μM SpbR and the stirring syringe was loaded with 120μL of 400μM RcdA. 19 injections of RcdA into SpbR were used to build the binding isotherm. Data analysis was performed with MicroCal (ORIGIN) and fitted to a single set of identical sites equation K_d_ = (Θ)/((1-Θ)~[X]), where Θ is the fraction of sites occupied by ligand X and [X] is the concentration of ligand X.

### Size-Exclusion Chromatography with Multi-Angle Light Scattering

Each protein complex was allowed to bind at room temperature for 45 minutes. The complexes were then injected onto a TSKgel™ G2000 or G3000 SEC column equilibrated in 20mM HEPES buffer, pH 7.0 with 100mM KCL (Tosoh Biosciences) at room temperature. The SEC column was coupled to an 18-angle light scattering detector (DAWN HELEOS-II) and a refractive index detector (Optilab T-rEX) (Wyatt Technology). Data was collected every second and the flow rate was set to 0.5mL/min. Data analysis was carried out using the program ASTRA (Wyatt Technology). Monomeric bovine serum albumin (BSA; Sigma) was used for calibration of the light scattering detectors and general data quality control. Measurements were taken at 25°C. The concentrations used in the SEC-MALS experiments are as follows unless otherwise noted in the figure legends: 25μM RcdA, 25μM RcdA L82E, 25μM RcdA 3E, 50μM PopA, 25μM SpbR, 15μM His-TacA.

### Microscale Thermophoresis (MST)

The MST experiments were performed using a Monolith NT.115 instrument (NanoTemper). Fluorescin-5-Maleimide labeled RcdA was incubated with increasing concentrations of unlabeled RcdA in the same buffer used in the polarization experiments. The measurements were performed at 20% MST power with 40% LED Power and with 3 s laser on time and 25 s off time. The K_d_ values were calculated using MO Affinity Analysis software (NanoTemper) and fit to the nonlinear equation Fnorm = [unbound + (bound-unbound) / 2 ~ (FluoConc + c + Kd – Sqrt((FluoConc + c + Kd)^2 – 4~FluoConc~c)], where unbound and bound are the thermophoresis values of the unbound and bound states, FluoConc is the fixed concentration of the fluorophore, Fnorm is the normalized fluorescence, and c is the concentration of the unlabeled protein.

### Hydrogen-Deuterium Exchange Mass Spectrometry

Hydrogen-deuterium exchange was measured on a Synapt G2Si high definition mass spectrometer (Waters). Deuterium exchange and quenching steps were performed using an automated HDX robotics platform (Waters). Samples were diluted 1:16 in D_2_O-containing buffer containing 0.1 mM K_2_HPO_4_ to final concentrations as specified in the figure legends. Deuterium exchange was allowed to take place for 0, 1, 10 and 60 minutes at 25°C, with staggered starts for each dilution reaction. After all reactions were completed, aliquots were removed and diluted 1:2 into cold quench buffer at 4°C (water with 4M Guanidine Hydrochloride at pH 2.5) and subsequentially run over an immobilized Water ENZYMATE immobilized pepsin column (ID: 2.1 length: 30mm) at a flow rate of 0.15mL/min at high pressure (~11000 psi) for peptide digestion. Prior to HDX analysis, the quality of each sample was assessed using SDS-PAGE and size-exclusion chromatography.

Three independently prepared experimental replicates and labeling reactions were performed for each condition and averaged in the peptide uptake plots. Blank runs were run in between each analysis to avoid peptide carry-over. Continuous lock-mass correction was performed using leu-enkephalin compound. Timepoints and analysis were randomized to ensure no biasing of results and to ensure variation. Peptides were ionized and separated by electrospray ionization for analysis in MSE mode at a mass resolution of 50-2000m/z range. Identification of peptides and analysis of the uptake plots and charge states for each peptide were completed in Protein Lynx Global Server (PLGS) and the software DynamX (Waters). Differential uptake heatmaps and uptake plots were plotted and created in Adobe Illustrator and in GraphPad Prism. Protections of greater than 15% are shown on the surface renditions of the RcdA structure (3CTW) using PyMol (Schrodinger Software)

### *In vitro* Degradation Assays

Degradation of proteins was monitored using SDS-PAGE gels as described previously (Bhat et al., 2013). The concentrations of different proteins used in degradation reactions are indicated in the figure legends. Degradation of GFP-CtrARD+15 was monitored with the loss of fluorescence over time as described previously (Smith et al., 2014). The final concentrations of each protein used in the reactions were as follows unless otherwise noted: 3μM RcdA, 3μM RcdA(L82E), 3μM RcdA(3E), 2μM CpdR, O.2μM CIpX, O.4μM CIpP, 1μM GFP-TacADBD, 4μM SpbR, 4μM TacA, 2μM GFP-CtrARD+15, 5mM ATP. 15μM RcdA(3E) was used in the GFP-CtrARD+15 degradation assay. GFP-CtrARD+15 curves were fit to a modified hyperbolic equation with the form of: Y = ((Vmax ~ [RcdA]) / (K_act_ + [RcdA])) + A, where A is a baseline constant, using GraphPad Prism.

### Microscopy

Phase contrast microscopy was performed on glass slides layered with a 1% agarose pad. A Zeiss Scope A.1 microscope (Zeiss, Germany) equipped with 100X (1×25 oil ∞/0.17) objective and 60 N-C 1” 100x camera was used. Images were analyzed with BacStalk (Drescher Lab, Max Planck Institute) software. Stalk distributions were compared using a one-tailed unpaired t-test with α = 0.05 (GraphPad Prism).

### Motility Assays

Motility assays were performed as described previously (Rood et al., 2012). Briefly, 0.3% agar plates containing varying concentrations of xylose and glucose were inoculated with three independent colonies of each strain for 3 days at 30°C. Colony sizes were determined using ImageJ (NIH). Quantifications were completed using ImageJ and plotted in GraphPad Prism.

### Sedimentation Velocity-Analytical Ultracentrifugation

Sedimentation Velocity experiments were completed using a Beckman ProteomeLab XL-1 Analytical Ultracentrifuge (Beckman). Samples were diluted into 20 mM HEPES pH 8.5, 100mM KCl, 10% glycerol at the concentrations indicated in the figure legends. The samples were spun at 55,000 g overnight at 25°C. The ρ and v values used for data fitting were determined using SEDNTERP and the amino acid sequence for each protein. The sedimentation velocity data was directly fit to the c(s) distribution method using the program SEDFIT and using the first 100 velocity scans for each condition. The resulting distributions from each experiment were then plotted in GraphPad Prism. The final concentrations used were the same concentrations used in the SEC-MALS experiments.

## Acknowledgements

We thank the Chien, Strieter, Stratton and Serio labs for helpful comments and discussions. We thank A. Kosowicz, W. Chowdhury and M. Sutherland for their prior work on the RcdA variants and generating plasmids. The anti-SpbR, anti-RcdA, and anti-FtsZ antibodies were graciously provided by G. Bowman, L. Shapiro, and E. Goley. This project was supported by funds from the NIH (R35GM130320). N.K. was supported in part through the Biotechnology Training Program (NIH T32GM108556). Special thanks to Lizz Bartlett in the IALS Biophysical Instrumentation Facility and Steve Eyles in the IALS Mass Spectrometry Facility for their technical assistance.

## Author Contributions

Conceptualization, N.K., P.C.; Investigation, N.K., D.D.; Writing, N.K., P.C.; Funding Acquisition, P.C.; Supervision, P.C.

