## Supplemental File for "Cargo competition for a dimerization interface stabilizes a protease adaptor in *Caulobacter crescentus*"

#### Supplemental Figure Legends

##### Supplemental Figure 1

**A)** Analytical ultracentrifugation of the SpbR-RcdA complex. A table of the measured masses is shown on the right. The concentrations for each component are the same as in the SEC-MALS experiments. **B)** Fluorescence polarization curve of the SpbR-RcdA complex. A serial dilution of unlabeled SpbR starting at 20μM was titrated into 200nM labeled RcdA. **C)** Fluorescence polarization curve of RcdA dimerization. A serial dilution of unlabeled RcdA starting at 100μM was titrated into 200nM labeled RcdA. **D)** Fluorescence polarization curve of the PopA-RcdA complex. A serial dilution of unlabeled PopA starting at 40μM was titrated into 200nM labeled RcdA.

##### Supplemental Figure 2

**A)** Uncropped *in vitro* gel-based degradation of RcdA alone or with SpbRDD. **B)** Uncropped *in vitro* gel-based degradation of RcdA in the absence or presence of excess RcdAΔC. **C)** Fluorescence polarization reporting on RcdA dimerization. 20μM unlabeled RcdA and RcdA L82E was added to each labeled reporter at 200nM. **D)** Uncropped *in vitro* degradation assay of RcdA and RcdA L82E. **E)** Chloramphenicol shutoff assays monitoring the degradation of RcdA and RcdA L82E in cells expressing wildtype RcdA or RcdA L82E. Quantification of the average of 3 biological replicates are shown at right.

##### Supplemental Figure 3

**A)** Fluorescence polarization curves of the PopA-RcdA or PopA-RcdAL82E complex. A serial dilution of unlabeled PopA starting at 40μM was titrated into 200nM labeled RcdA (same as shown in Supplemental Figure 1) or 200 nM labeled RcdA L82E. **B)** Uncropped *in vitro* degradation assay of SpbR with RcdA and RcdA L82E. **C)** Uncropped *in vitro* degradation assay of TacA with RcdA and RcdA L82E. **D)** GFP-CtrA reporter degradation curves with RcdA and RcdA L82E. Fits are to a modified hyperbolic equation as outlined in the methods. **E)** SEC-MALS traces of the L82E-PopA complex. 50μM PopA was incubated with 50μM RcdA L82E.

**Supplemental Figure 4**

**A)** Differential uptake heatmap plot of 20μM versus 600nM RcdA to illustration dimerization interface. Given $K_D$ determined in Figure 1, 20 uM would be primarily dimer while 600 nM would be primarily monomer. **B)** Crystallographic dimer interface as predicted by the crystal structure and PDBePISA interface predictions (PDB ID: 3CTW). **C)** Individual deuterium uptake plots for regions displaying >15% protection in the RcdA-L82E, RcdA-$^{DBD}$TacA, and RcdA-PopA HDX datasets. Quantification of three independent experimental replicates is shown with the mean and standard deviation plotted as error bars.

**Supplemental Figure 5**

**A)** Sequence alignment of RcdA in other alpha-proteobacteria (WebLogo, Berkeley). The upper panel highlights in red the residues (R49, K53, R57) mutated to glutamic acid to general RcdA 3E. The lower panel highlights in red the L82 residue mutated to glutamic acid to make RcdA L82E. **B)** Uncropped *In vitro* degradation assay of SpbR with RcdA and RcdA 3E. **C)** Uncropped *In vitro* degradation assay of TacA with RcdA and RcdA 3E.

**Supplemental Figure 6**

**A)** Western blot of steady state levels of integration strains expressing RcdA, L82E or 3E from the xylose locus at low induction (0.002% xylose). Quantifications are normalized to wildtype levels. **B)** Quantifications of 3 biological replicates of the motility assays in low induction (+xylose) and no induction (+glucose). **C)** Uncropped western blots of the SpbR and CtrA shutoffs for the L82E mutant. **D)** Uncropped western blots of the SpbR and CtrA shutoffs for the 3E mutant.

### Supplementary Figure 1

A

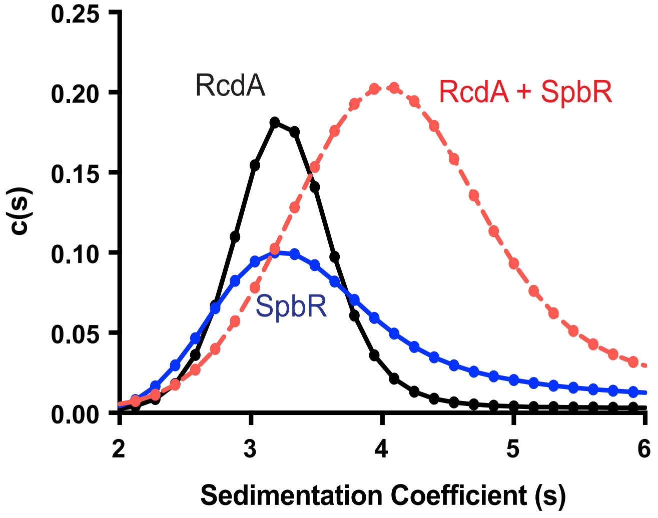


B

| Protein | Sedimentation Coefficient (s) | Experimental Mass (kDa) |
|---|---|---|
| RcdA | 3.0 ± 0.06 | 37.0 |
| SpbR | 3.0 ± 0.03 | 41.0 |
| RcdA + SpbR | 4.2 ± 0.05 | 60.1 |

C

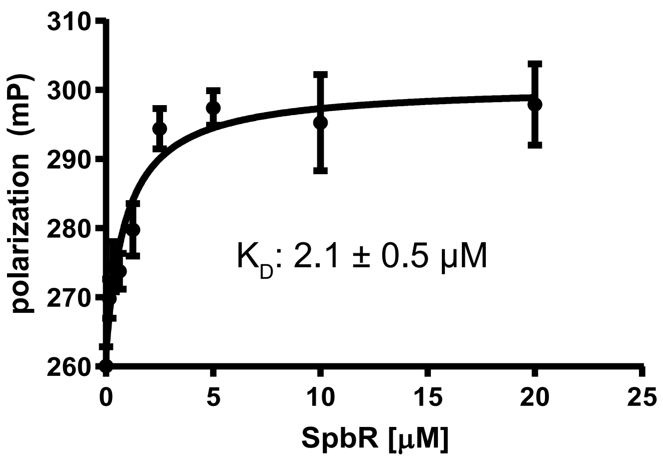


D

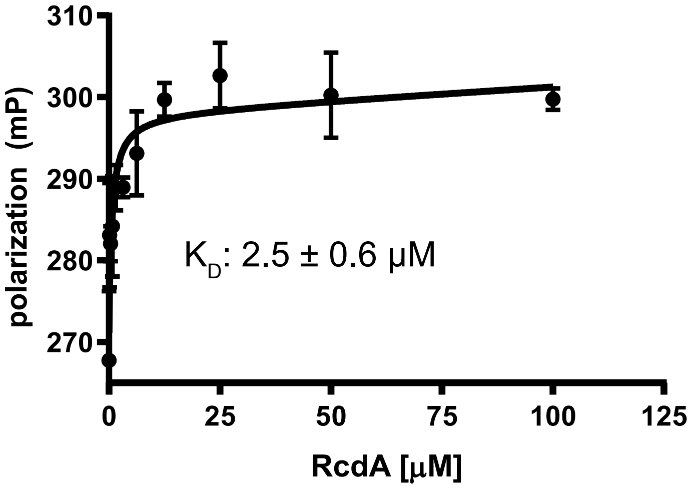


E

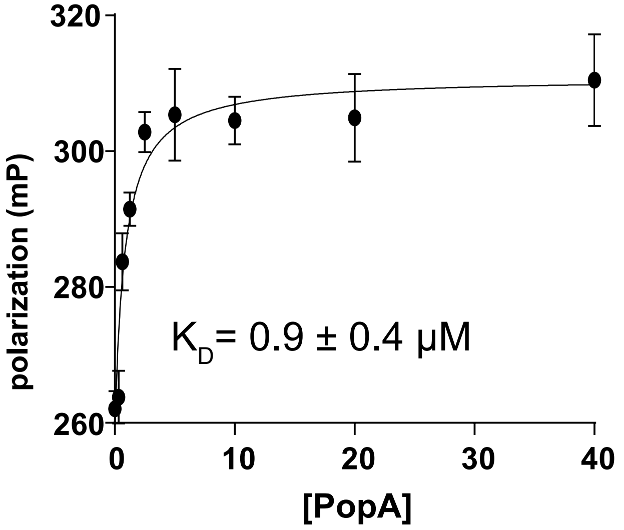

### Supplementary Figure 2

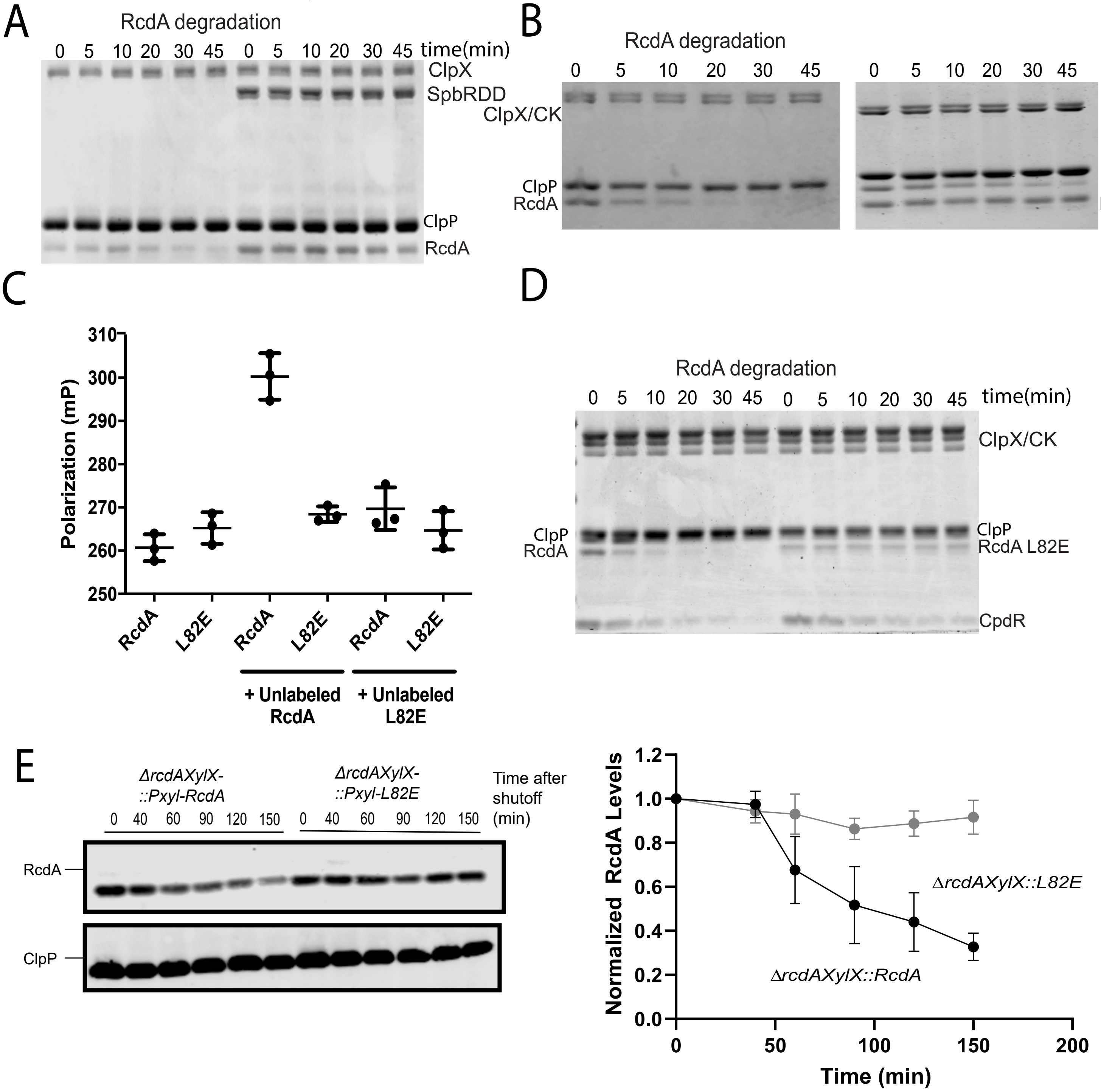

### Supplementary Figure 3

A

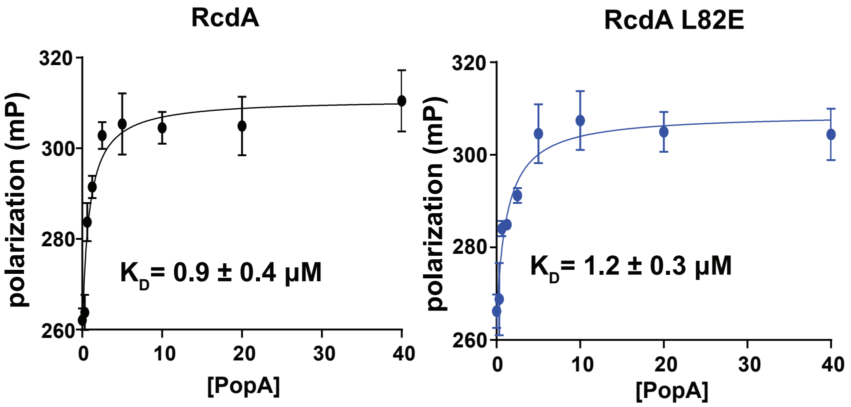


B

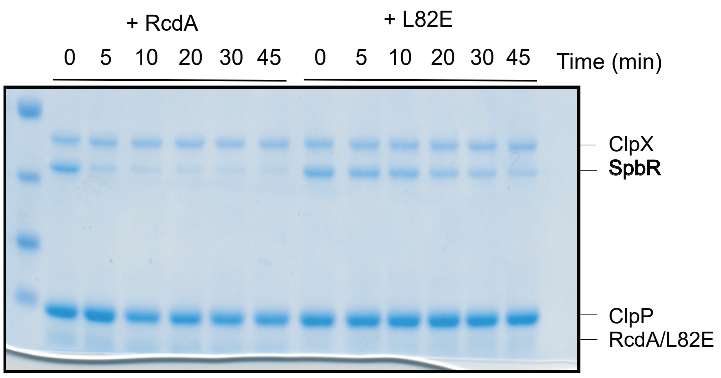


C

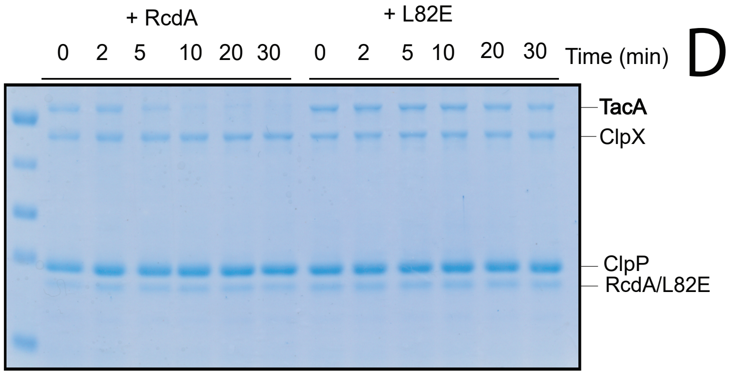


D

| Protein | $K_{act}$ (μM) | Vmax (GFP-CtrARD+15/ClpX$_6$/Min) |
|---|---|---|
| RcdA | 0.22±0.09 | 0.04±0.04 |
| L82E | 0.11±0.05 | 0.04±0.05 |

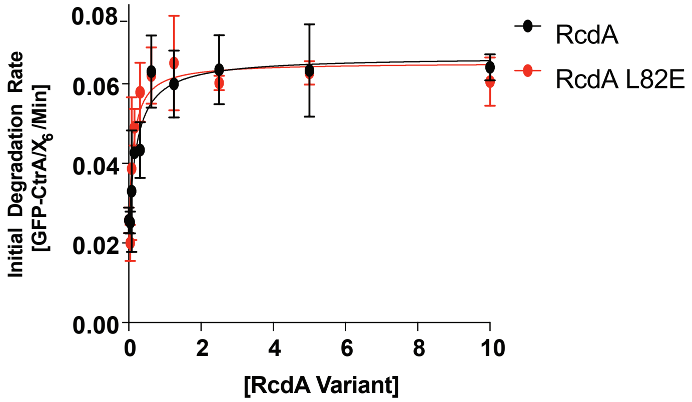


E

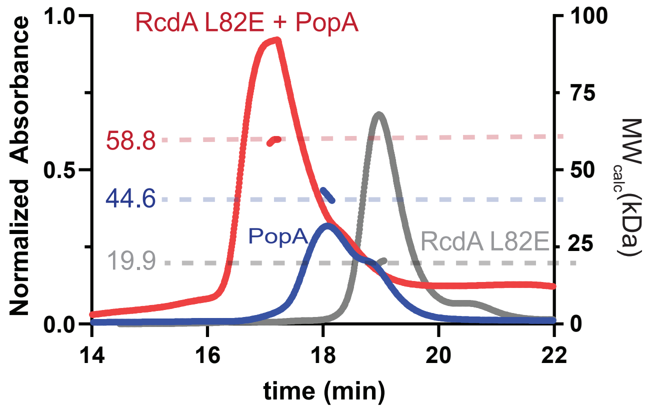

### Supplementary Figure 4

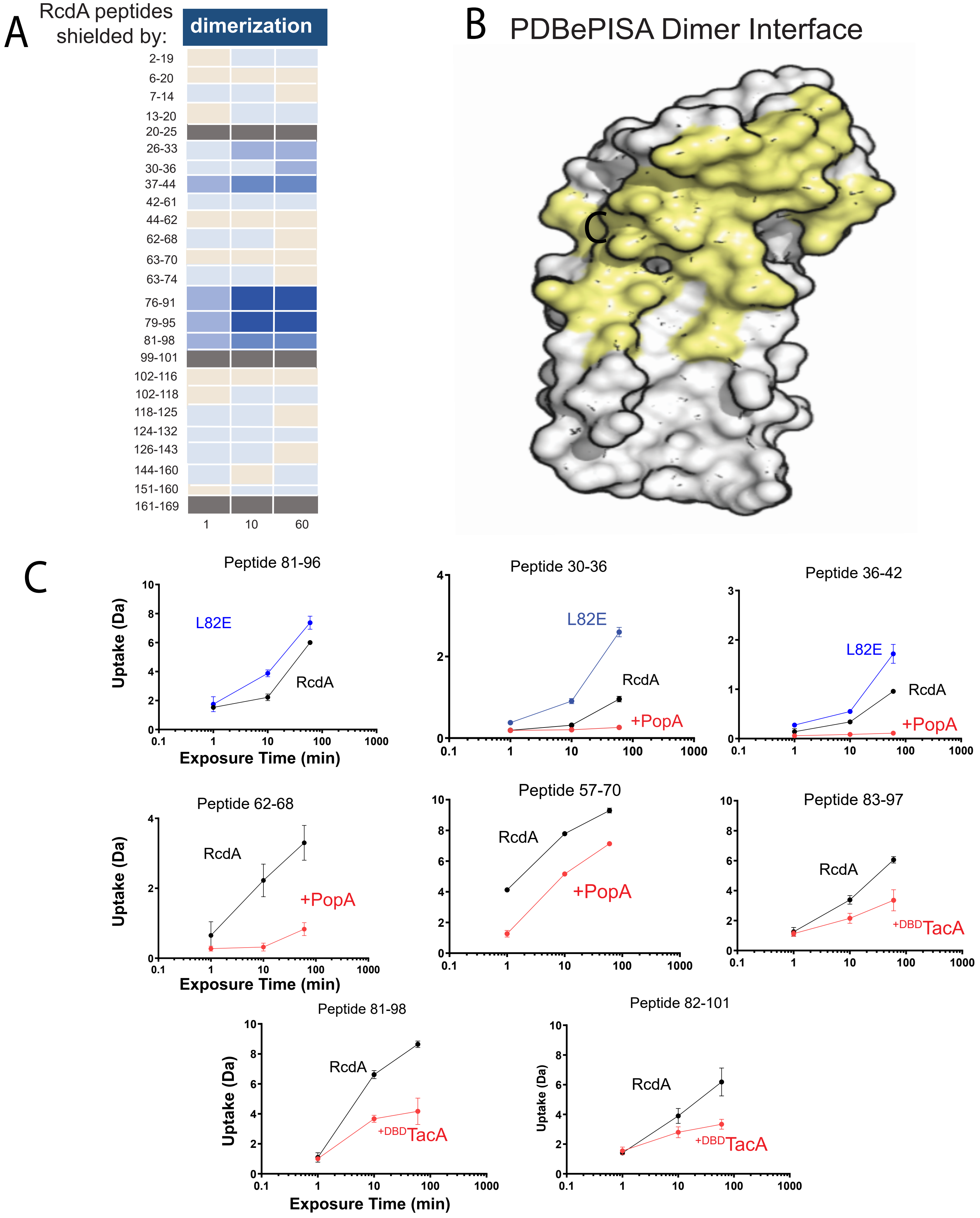

### Supplementary Figure 5

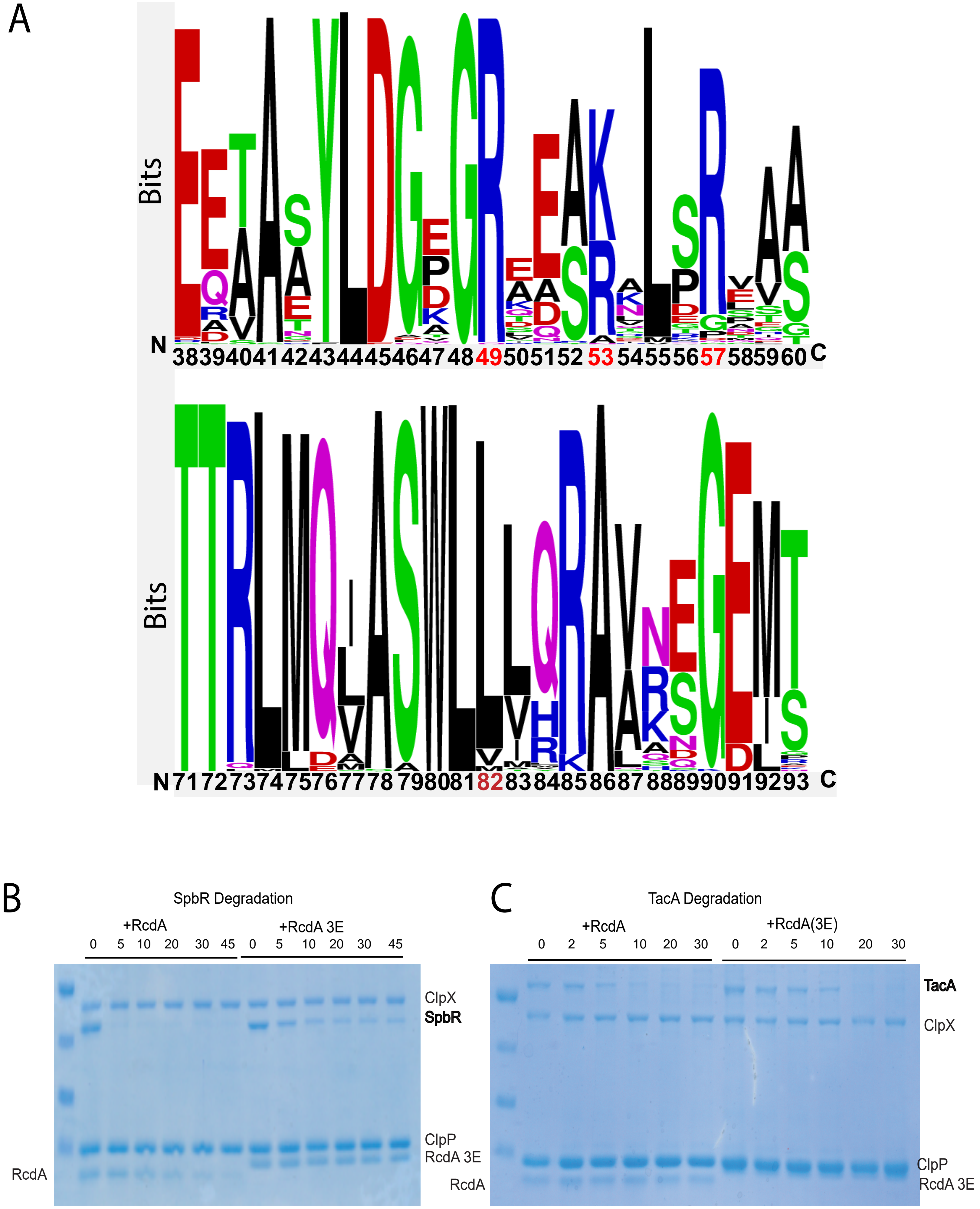

### Supplemental Figure 6

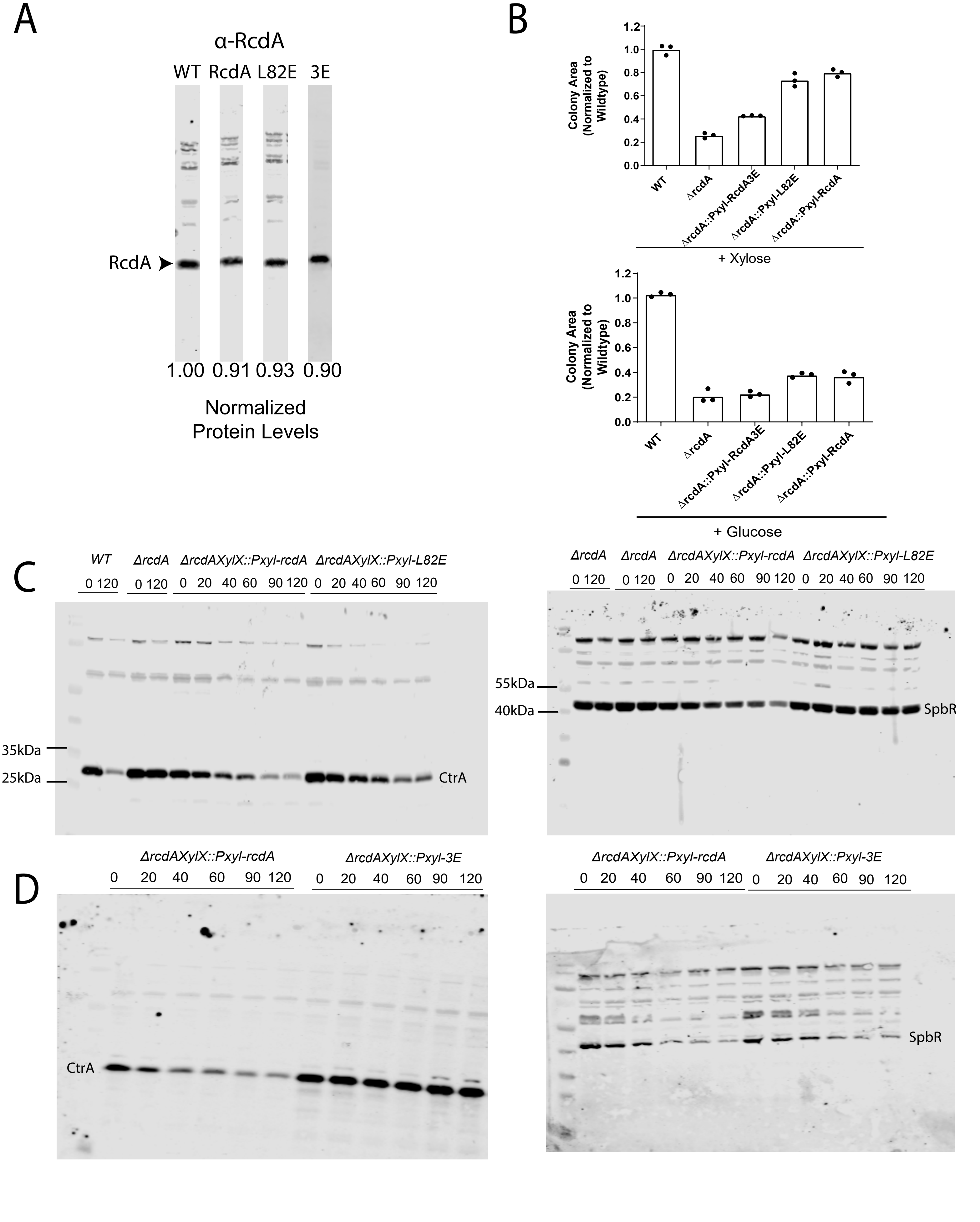

### Supplementary Table 1

| Organism | Name | Description | Source |
|---|---|---|---|
| *C.crescentus* | CB15N | synchronizable derivative of wild-type CB15 | (Evinger and Agabian, 1977) |
| | CPC452 | *ΔrcdA (hyg$^{R}$)* | (Mcgrath et al., 2006) |
| | CPC963 | *ΔrcdAXylX::Pxyl-RcdA* | This study |
| | CPC964 | *ΔrcdAXylX::Pxyl-L82E* | This study |
| | CPC965 | *ΔrcdAXylX::Pxyl-3E* | This study |
| *E. coli* | TOP10 | Cloning strain | Invitrogen |
| | BL21(DE3) pLysS | recombinant protein expression | Invitrogen |
| | X90 T7 | recombinant protein expression | From Bob Sauer |
| | EPC100 | dh5alpha pQE70-his-ClpP | (Chien et al., 2007) |
| | EPC112 | BL21DE3 pLysS pET23 Ulp1his protease | This study |
| | EPC162 | BL21DE3 375 eGFP-His6-CtrARD+15 | (Smith et al., 2014) |
| | BPC180 | Top10 pXMCS-2 | This study |
| | BPC238 | BL21DE3 plysS pET23 ClpX | (Chien et al., 2007) |
| | EPC196 | BL21DE3 plysS pET23b His6Sumo-CpdR | (Lau et al., 2015) |
| | EPC677 | BL21DE3 plysS 375 His6-TacA | (Joshi et al., 2015) |
| | EPC811 | BL21DE3 plysS pET23b His6Sumo-TacADBD (437-488) | (Joshi et al., 2015) |
| | EPC812 | BL21DE3 plysS pET23b His6Sumo-SpbR | (Joshi et al., 2015) |
| | EPC960 | BL21DE3 plysS pET23b His6Sumo-SpbRDD | (Joshi et al., 2015) |
| | EPC967 | BL21DE3 plysS pET23b His6Sumo-RcdAΔC | (Joshi et al., 2015) |

| | | | |
|---|---|---|---|
| | EPC970 | BL21DE3 plysS pET23b His6Sumo-RcdA | (Joshi et al., 2015) |
| | EPC1000 | TOP10 pET23b His6Sumo-RcdA | (Joshi et al., 2015) |
| | EPC1037 | BL21DE3 plysS pET23b His6Sumo-PopA | (Joshi et al., 2015) |
| | EPC1129 | BL21DE3 plysS pET23b His6Sumo-RcdA 3E | This study |
| | EPC1517 | BL21DE3 plysS pET23b His6Sumo-RcdA L82E | This study |
